# The recombination landscape of *Drosophila melanogaster* can be repatterned by a single gene

**DOI:** 10.1101/2025.05.16.654569

**Authors:** Stacie E. Hughes, Cynthia Staber, Grace McKown, Zulin Yu, Justin P. Blumenstiel, R. Scott Hawley

**Affiliations:** Stowers Institute for Medical Research, Kansas City, Missouri, 64110, United States of America; Department of Ecology and Evolutionary Biology, University of Kansas, Lawrence, Kansas, 66045, United States of America

**Keywords:** meiosis, crossover, synaptonemal complex Running title: *D*. *mauritiana* C(3)G alters the crossover pattern

## Abstract

Meiotic recombination plays an important role in ensuring proper chromosome segregation during meiosis I through the creation of chiasmata that connect homologous chromosomes. Recombination plays an additional role in evolution by creating new allelic combinations. Organisms display species-specific crossover patterns, but how these patterns are established is poorly understood. *Drosophila mauritana* displays a differing recombination pattern compared to *Drosophila melanogaster*, with *D. mauritiana* experiencing a reduced centromere effect, the suppression of recombination emanating from the centromeres. To evaluate the contribution of the synaptonemal complex (SC) C(3)G protein to these recombination rate differences, the *D. melanogaster* allele was replaced with *D. mauritiana c(3)G* coding sequence. We found that the *D. mauritiana* C(3)G could interact with the *D. melanogaster* SC machinery to build full length tripartite SC and chromosomes segregated accurately, indicating sufficient crossovers were generated. However, the placement of crossovers was altered, displaying an increase in frequency of the centromere-proximal euchromatin indicating a decrease in the centromere effect similar to that observed in *D. mauritiana*. Recovery of chromatids with more than one crossover was also increased, likely due to the larger chromosome span now available for crossovers. As replacement of a single gene mediated a strong shift of one species’ crossover pattern towards another species, it indicates a small number of discrete factors may have major influence on species-specific crossover patterning. Additionally, it demonstrates the SC, a structure known to be required for crossover formation in many species, is likely one of these discrete factors.

**Lay Abstract:** Meiotic crossovers are important for ensuring proper chromosome segregation and generating genetic diversity. Different species display unique crossover patterns but the mechanisms that establish these patterns are poorly understood. The synaptonemal complex (SC) is built between meiotic chromosomes and promotes crossover formation. Replacement of the SC gene *c(3)G* in the fruit fly *Drosophila melanogaster* with *Drosophila mauritiana c(3)G* resulted in full-length SC assembly and proper chromosome segregation, but the *D. melanogaster* crossover pattern was shifted to appear more similar to *D. mauritiana*. This demonstrates that crossover patterning can be largely influenced by minor changes in the makeup of the SC.

## Introduction

Meiosis is a specialized cell division that results in haploid gametes. In many organisms, there are several important steps that must occur for meiosis to be successful. Homologous chromosomes pair, double-strand breaks (DSBs) are induced, chromosomes undergo synapsis, and a subset of the DSBs are repaired into crossovers that physically link the homologous chromosomes throughout prophase. These physical linkages, known as chiasmata, act in conjunction with cohesion between the sister chromatids and spindle microtubules to ensure homologs properly orient and segregate towards opposite spindle poles during the meiosis I division. Failure to generate a crossover on each chromosome can lead to the missegregation of the homologous chromosomes and the production of aneuploid gametes that cause miscarriage, birth defects, and infertility (Hassold and Hunt 2001; Nagaoka *et al*. 2012).

While a single crossover per chromosome can ensure proper segregation at the meiosis I division, many organisms make more than one crossover per chromosome. For example, *Drosophila melanogaster* females make an average of one crossover per chromosome arm, which results in two crossovers per autosome, and yeasts, such as *Saccharomyces cerevisiae,* make multiple crossovers per chromosome (Dutta *et al*. 2024; Lindsley and Sandler 1977; Miller *et al*. 2016). The honey bee, *Apis mellifera*, has a very high level of meiotic recombination, with more than five recombination events per chromosome pair per meiosis (Beye *et al*. 2006). Additionally, organisms differ in how crossovers are distributed across the chromosomes with some chromosomal regions more or less likely to experience a crossover. For example, in many mammals, recombination preferentially occurs near “recombination hotspots” that are dependent on PRDM9 (reviewed in (Grey *et al*. 2018). Conversely, recombination is suppressed in the repetitive heterochromatin in most species (Fernandes *et al*. 2019; Pazhayam *et al*. 2025; Pazhayam *et al*. 2021; Smith and Nambiar 2020; Topp and Dawe 2006). These differences in crossover number and placement lead to species-specific recombination patterns that can differ even between relatively closely related species (Dumont 2017; Dutta *et al*. 2024; Stapley *et al*. 2017; True *et al*. 1996). These differences can potentially influence evolution because recombination rate and placement affect the likelihood that alleles will be inherited together in offspring. Regions of low recombination may accumulate deleterious genetic elements, such as transposable elements, due to Muller’s Ratchet, which is the accumulation of deleterious elements due to the stochastic loss of the most fit haplotype (Dolgin and Charlesworth 2008). Recombination, by uncoupling linked alleles, provides an opportunity to eliminate, from a population, less fit alleles that are linked to beneficial alleles residing on the same chromosome (reviewed in (Comeron *et al*. 2008; Rice 2002)). Since crossover rate and placement can affect the capacity of a species to adapt to changing conditions, it is important to understand the factors that can influence crossover patterning.

The fruit fly *Drosophila melanogaster* has long been a model for studying aspects of crossover patterning, such as crossover interference (the designation of one crossover inhibiting the designation of another crossover nearby) and the centromere effect (Lindsley and Sandler 1977; Sturtevant 1913; Sturtevant 1915). The centromere effect is the polar suppression of recombination emanating from the centromeres (reviewed in (Hartmann *et al*. 2019; Pazhayam *et al*. 2024; Pazhayam *et al*. 2021). The centromeres in *D. melanogaster* are flanked by large blocks of heterochromatin which are also regions where recombination is suppressed. The centromere-proximal heterochromatin is thought to act as a buffer, functioning as an inert spacer, against the centromere effect in *D. melanogater* rather than directly providing the mechanism for the crossover suppression in the centromere-proximal euchromatin. In experiments that move centromere proximal heterochromatin away from the centromere, thus bringing euchromatic sequences closer to the centromere, those centromere-proximal euchromatic regions experience increased levels of crossover suppression (Schalet and Lefevre 1976; Yamamoto and Miklos 1977; Yamamoto and Miklos 1978). These experiments indicate the centromere effect is mediated by the centromere itself or trans-acting factors rather than the surrounding heterochromatin. While these experiments ruled out the role of the heterochromatin in mediating the centromere effect, they were not designed to distiniguish whether the centromere effect was mediated by *trans*-acting factors, *cis*-acting elements within the centromere, or a combination of the two.

Interestingly, both True *et al*. (1996) and Hawley *et al*. (2025) found that the centromere effect was lessened in the closely related species *Drosophila mauritiana*. An increase in crossing-over was observed in euchromatic regions proximal to the centromeres in *D. mauritiana* compared to *D. melanogaster* (Hawley *et al*. 2025; True *et al*. 1996). Likely due to the decreased centromere effect increasing the span available to acquire a crossover on most chromosome arms, double crossover chromatids and total map length, in *D. mauritiana* were increased (Hawley *et al*. 2025; True *et al*. 1996).

The mechanisms controlling crossover patterning have only just begun to be elucidated and even less understood is how species-specific patterns evolve. Are changes in crossover patterning the result of small changes in many genetic loci, or can global patterning be shifted by changes in just a few genes? One *trans-*acting factor that appears to mediate the differences in crossover patterning between *D. melanogaster* and *D. mauritiana* is Mei-218 (Brand *et al*. 2018). Mei-218 is a pro-crossover protein and part of the mei-MCM complex (Kohl *et al*. 2012). When the *D. mauritiana* version of *mei-218* was expressed as a transgene in *D. melanogaster* females, in the absence of a functional endogenous *mei-218* locus, the crossover pattern of the *D. melanogaster* flies displayed an increase in total crossing over compared to a *D. melanogaster mei-218* transgene (Brand *et al*. 2018). However, both transgenes displayed a pattern that was somewhat distinct from wild type. As crossover patterns are complex there may be other factors that majorly influence crossover patterning.

A meiotic structure that may influence meiotic crossover patterning is the synaptonemal complex (SC). The SC is a ladder-like structure that builds along the homologous chromosome during early pachytene. In many species, the SC is required for crossover formation (reviewed (Page and Hawley 2004; Zickler and Kleckner 2015)). While the physical structure of SC appears conserved between divergent species based on electron microscopy, the individual proteins of the SC are rapidly evolving at the amino acid level. SC proteins can be difficult to identify using homology to amino acid sequence alone (Hemmer and Blumenstiel 2016; Kursel *et al*. 2021; Williams and Hawley 2025; Zakerzade *et al*. 2025). Because the SC proteins are rapidly evolving, they have the potential for being another avenue for regulating crossover patterning between species. It has been shown previously that conditions that perturb the SC can result in changes in crossover number and location (Billmyre *et al*. 2019; Kohler *et al*. 2025; Libuda *et al*. 2013; Pazhayam *et al*. 2025). For example, in *C. elegans,* partial depletion of a SC component led to increases in COSA-1 foci, which mark crossovers, and a decrease in crossover interference (Libuda *et al*. 2013). In *D. melanogaster,* in-frame deletions within the coiled-coil domain of the transverse protein C(3)G resulted in SC that disassembled earlier than wild type, along with a shift of crossovers towards the centromere-proximal euchromatin (Billmyre *et al*. 2019). In *Arabidopsis thaliana,* the SC appears non-essential for the formation of crossovers, but loss of the SC leads to increased crossover formation, suggesting the role of the SC is regulating crossover interference in this species (Capilla-Perez *et al*. 2021).

To more directly test the role of SC components in crossover patterning and the impact that SC sequence divergence has on recombination, we tested whether the replacement of the *D. melanogaster* SC protein C(3)G with the *D. mauritiana* version of C(3)G could change crossover patterning. C(3)G is the *Drosophila* major transverse filament protein that spans across the SC and is the functional ortholog to Zip1 in yeast and SCP1 in mammals (reviewed (Gao and Colaiacovo 2018; Page and Hawley 2004)). The *D. melanogaster* C(3)G protein shares 89.1% amino acid identity with the *D. mauritiana* C(3)G protein. We show that replacing the *D. melanogaster c(3)G* with the *D. mauritiana* version changes the crossover patterning of *D. melanogaster* to more closely resemble the *D. mauritiana* pattern, with an increase of crossovers in the centromere-proximal euchromatin and an increase in the recovery of double crossover chromatids. These results suggest that the difference in crossover patterning between *D. melanogaster* and *D. mauritiana* and the centromere effect in *D. melanogaster* are largely regulated by a small number of *trans-*acting factors, including C(3)G. In addition, our findings demonstrate that the rapid evolution of the SC proteins is likely to have functional consequences for establishing crossover patterning.

## Methods

### Drosophila husbandry

Flies were maintained at 25°C on standard food. c*(3)G^mau^* genotype is *y w/ y+Y; (3)G^mau^; sv^spa^-^pol^*except for recombination experiments (see Recombination methods). Control (c(3)G^+^) stock was *y w/ y+Y; sv^spa^-^pol^* to be genetically similar to the c*(3)G^mau^*stock, with the exception of recombination experiments (see Recombination methods).

The replacement of the *c(3)G* gene in *D. melanogaster* was performed by WellGenetics using CRISPR. A donor template containing a *D. melanogaster* codon optimized coding region of the *D. mauritiana c(3)G* was constructed. Codon optimization was performed using the IDT DNA codon optimization tool (https://www.idtdna.com/CodonOpt) with the gBlocks option (Supplementary Fig. 1). Codon optimization was used to ensure robust expression of the *D. mauritiana c(3)G* gene, and by varying many of the synonymous positions, to limit difficulties of homologous replacement using CRISPR. CRISPR-mediated mutagenesis for the codon-optimized *D. mauritiana c(3)G* replacement was performed by WellGenetics Inc. using modified methods described in Kondo and Ueda (2013). Briefly, the upstream gRNA sequence ACGGGCGCCAAGAAAAGATC[GGG] and the downstream gRNA sequence CTTGATGGGCGCACATAATT[TGG] were cloned into U6 promoter plasmids. The cassette Dmau c(3)G cDNA-PBacDsRed, containing 3xP3-DsRed flanked by PiggyBac terminal repeats, and two homology arms were cloned into pUC57-Kan as donor template for repair. *c(3)G/CG17604*-targeting gRNAs and hs-Cas9 were supplied in DNA plasmids, together with donor plasmid, for microinjection into embryos of control strain *[SWGa3411] w1118 hyb5b-8*. F1 flies carrying the 3xP3-DsRed selection marker were further validated by genomic PCR and sequencing. The CRISPR generated a 2,655-bp deletion of *c(3)G/CG17604,* which deleted the entire CDS of *c(3)G/CG17604* that was replaced by cassette Dmau c(3)G cDNA-PBacDsRed. The 3xP3-DsRed cassette was then excised through PiggyBac transposition, and the excision was validated by PCR and sequencing.

The *c(3)G^mau^*gene was integrated into the *y w/ y+Y; sv^spa^-^pol^* genetic background using standard genetic crosses and the replacement PCR verified.

### Immunofluorescence

For immunofluorescence of whole-mount ovaries, females 1-3 days post-eclosion were provided with wet yeast paste and males overnight at 25°C. Ovaries were dissected in PBS plus 0.1% Tween-20 (PBST). Ovaries were fixed for 20 min in 200 µL of PBS containing 2% formaldehyde (Ted Pella) and 0.5% Nonidet P-40 plus 600 µL heptane. After three washes in PBST, ovaries were blocked for at least one hour in PBST plus 1% bovine serum albumin (BSA) (EMD Chemicals, San Diego, CA). For experiments using the γH2Av antibody, the primary antibodies were applied in PBST plus 0.5% BSA overnight at 4°C, while for all other experiments primary antibodies were applied in PBST overnight. For experiments using the γH2Av antibody, ovaries were washed three times in PBST before blocking again for at least one hour in PBST plus 1% BSA. Secondary antibodies were applied overnight at 4°C in PBST plus 0.5% BSA. For all other experiments, after the three washes in PBST, secondary antibodies were applied for 3-5 hours at room temperature. For all experiments, 1.0 µg/ mL DAPI was applied during the last 15 minutes of secondary antibody incubation. Samples were washed three times in PBST. Ovaries were mounted in Prolong Gold or Prolong Glass (Thermofisher).

Primary antibodies rat anti-*D. mauritiana* C(3)G (see Antibody Production methods), rabbit anti-histone γH2AVD pS137 [1:5000] (Rockland Inc., Lot 46042), mouse anti-Orb 6HA [1:40] (Developmental Studies Hybridoma Bank, Iowa) (Lantz *et al*. 1994), rabbit anti-Corolla [1:2000] (Collins *et al*. 2014), rabbit anti-Centromere Identifier (CID) [1:2000] (gift of Dr. Gregory Rogers), and mouse anti-lamin Dm0 ADL84.12 [1:100] (Developmental Studies Hybridoma Bank, Iowa) were used. The following goat Alexa Fluor secondary antibodies at [1:500] were used: anti-mouse 647, anti-rat 488, 555, and 594, and anti-rabbit 488 or 555 (ThermoFisher).

### Microscopes and imaging conditions

For most images, a Deltavision Elite system from GE Healthcare equipped with an Olympus IX70 inverted microscope and a high-resolution CCD camera was used. SoftWoRx v. 7.2.1 (Applied Precision/Leica Microsystems, Inc,) was used to deconvolve images. SoftWoRx v. 7.2.1 or Fiji were used for image analysis. Brightness and contrast were adjusted minimally for improved publication quality.

The STED imaging in Fig. 2 was performed using a Leica SP8 Gated STED Microscope equipped with a 100×, 1.4 NA oil immersion objective. The C3G channel (labeled with Alexa 594) was excited using a pulsed white light source (80 MHz) set to 594 nm, in combination with a pulsed 775 nm laser operating at 80–90% of its maximum output. All STED images were captured in 2D mode to optimize lateral resolution, with each image averaged eight times in line average mode. Emission photons were detected using internal Leica HyD hybrid detectors, with a time gate setting of 1–6 ns. Raw STED images were subsequently deconvolved using Huygens Professional software (version 14.10; Scientific Volume Imaging), employing a theoretically estimated point spread function for the deconvolution process. Default system parameters were used, except for the background, which was measured directly from the raw images, and the signal-to-noise ratio, which was adjusted to a range of 15–20.

**Figure 1.**
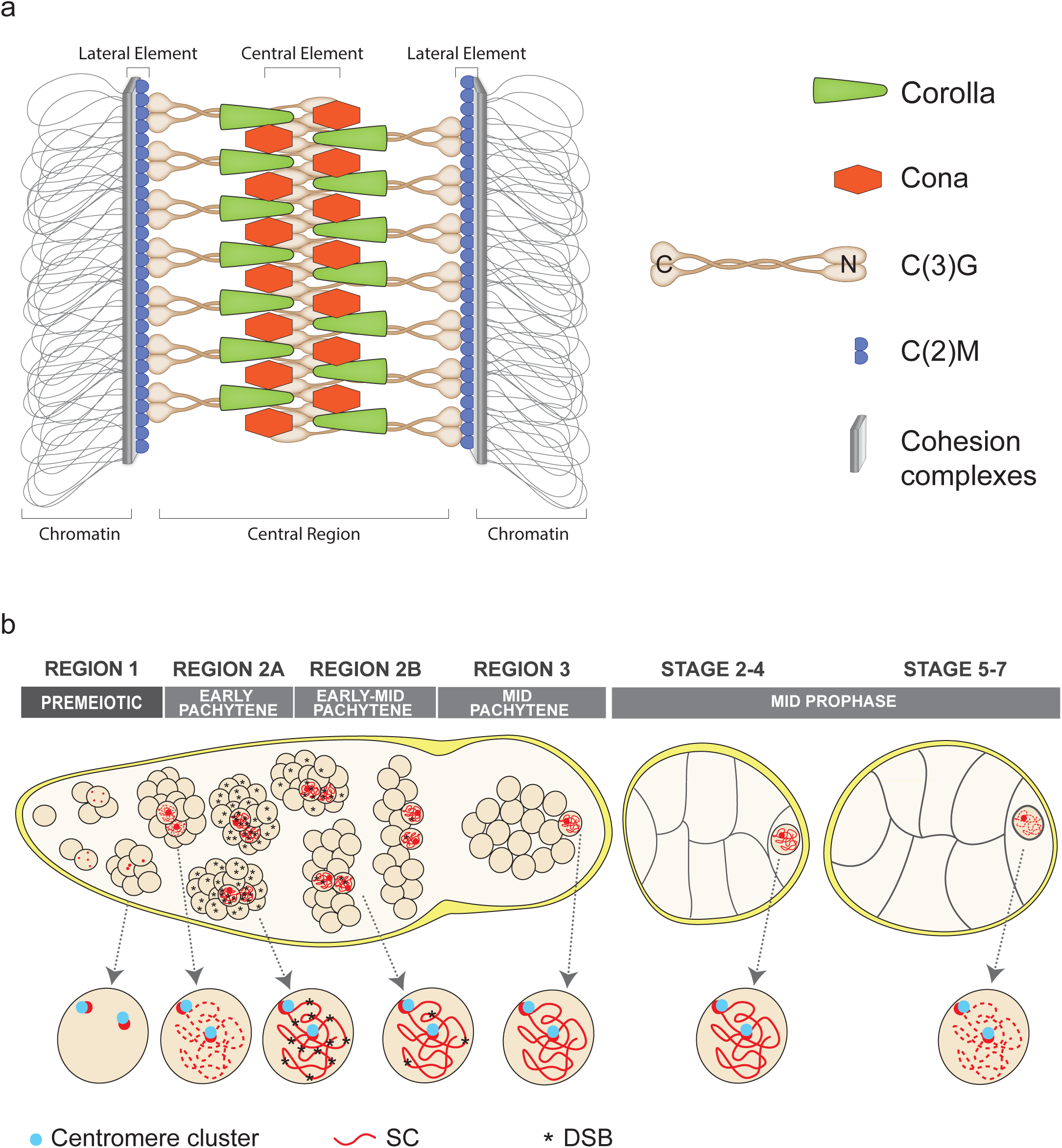
a) Model of the *D. melanogaster* SC. Cohesin complexes (represented as a gray bar) interact with the homologous chromosomes. The protein C(2)M (blue partial circles) is present along the chromosomes at the LE. Dimers of the transverse filament protein C(3)G (brown) in the central region connect to the LEs at the C-terminal end of the dimers and the N-terminal part of the dimers interacting with opposing C(3)G dimers. C(3)G is stabilized by the proteins Cona (orange hexagons) and Corolla (Green triangles) in the central element. b) Diagram of the meiotic events during the early developmental stages of the ovary. Individual nuclei are shown enlarged below the illustration of development. In the premeiotic region 1, the cystoblast divides four times to make a 16-cell interconnected cyst. SC proteins (red) begin to load near the centromeres (blue circles in the enlarged nuclei) that begin to cluster during the mitotic divisions. At the entry to region 2A (early pachytene), SC components load at additional sites along the chromosome arms. In region 2A the SC is full-length in 2-4 nuclei of each cyst, DSBs (asterisks) are induced in the cysts, and the centromeres are clustered into an average of two clusters. As the cysts progress to region 2B (early-mid pachytene), DSBs become progressively repaired based on γH2Av and two SC-positive nuclei are present within the cysts. By region 3 (mid-pachytene) the pro-oocyte is the only nucleus with full-length SC and the γH2AV marker for DSBs is absent. Centromeres remained clustered to an average of two foci. After the egg chamber buds off from the germarium the SC progressively disassembles as the egg chamber continues development.

**Figure 2.**
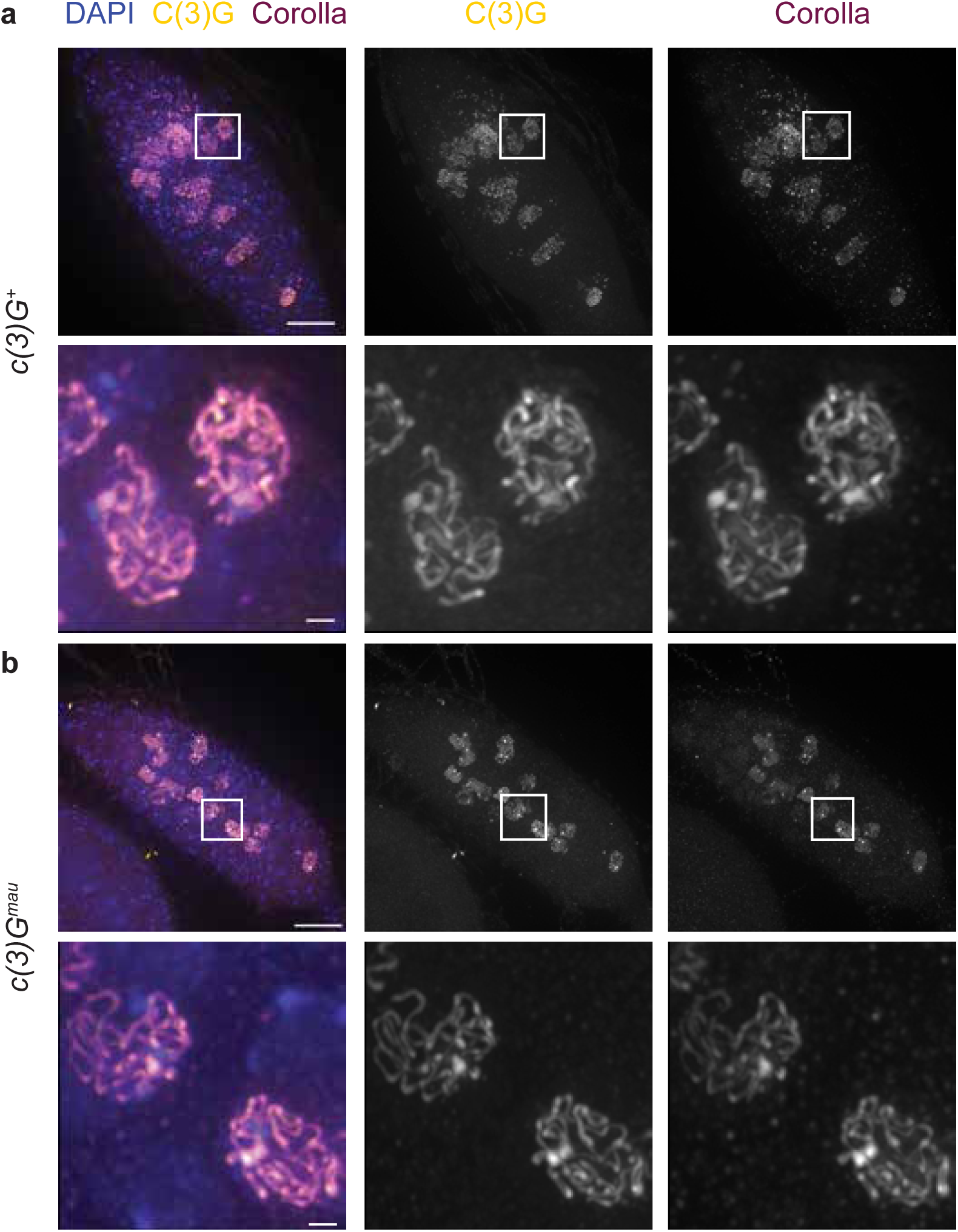
Full length SC assembles in *c(3)G^mau^* ovaries. a-b) Shown are antibodies raised against *D. melanogaster* Corolla (magenta) and the C-terminal end of *D. mauritiana* C(3)G (mau C(3)G) (yellow). DAPI is shown in blue. Boxed regions are shown enlarged. a) In *c(3)G^+^* the mau C(3)G antibody appears as full-length tracks that overlap with the Corolla immunofluorescence indicating the mau C(3)G antibody recognizes *D. melanogaster* C(3)G. b) In *c(3)G^mau^* both mau C(3)G and Corolla antibodies appear as full-length tracks in pachytene similar to *c(3)G^+^* indicating SC assembly occurs normally in *c(3)G^mau^*. Scale bar = 10 µm in germaria and 1 µm in enlarged images. Images are projections from z-stacks.

To measure the distance between tracks of the C-terminal mau C(3)G antibody in STED images, line profile measurements were acquired from line ROIs drawn manually perpendicularly across regions of the SC that appeared to be flat in the z dimension.

The average and standard deviation for the measurements were calculated for the measurements in Excel and a Mann-Whitney test was used for statistical comparisons between the distances for *c(3)G^+^* and *c(3)G^mau^* .

### Antibody Production

For the production of the *D. mauritiana* C(3)G antibody a construct encoding 324 amino acids in the C-terminal part of the protein (NESYLNELTETKLKHTQEIKEQADAYEIVVQELKESLNKASVDFTQLKSNSEKLHKETLLQVSQLQEKLTEMVSHRSNQEELVKRLGDELQEKTHSFEEELKRQQVQLANQMQMKTT EVASENKRNAVQIQTLKSELEDRNKAFKAQQDKLEFLISDLDKLKNAIINLQAEKMEIESELSTAKVKFSEELQFQKDSLMKKVSELELEIKRKENELIELEREKNNEMAVLQFKMNRINCVIDQPVTTIKQLKDSKGPTKTESIPTKVQPENTDGFSGTAKKRNARRQKITTYSSDFDSGDDMPILGNSLNGKRVKLCTPIKPQNN) (See Supplementary Fig. 2) with a start codon and N-terminal His tag was made in a pET30a by GenScript USA, Inc. The vector was expressed in BL21 (DE3) bacterial cells, and the protein was purified using Ni-NTA His-Bind Resin (EMD Millipore Corp) under denaturing conditions. The purified protein was used to generate an antibody in rats by Cocalico Biologicals, Inc.

### Recombination

For recombination studies of the *X* chromosomes virgin *y^1^ sc^1^ cv^1^ v^1^ f^1^ y+/y w; sv^spa^-^pol^/+* (*c(3)G^+^*) or *y^1^ sc^1^ cv^1^ v^1^ f^1^ y+/y w; c(3)G^mau^* ; sv*^spa-pol^/+* (*c(3)G^mau^*) females were crossed individually to *y sc cv v f car^1^/ B^S^Y* males. Only female progeny were scored for the genetic markers.

For *3^rd^* chromosome recombination scoring a recombinant *ru hry Diap1^1^ (th) st cu c(3)G^mau^* stock was constructed using standard genetic methods. Virgin *y w/ +; ru^1^ hry^1^ Diap1^1^(th) st^1^ cu^1^ sr^1^ e^s^ ca^1^ /+; sv^spa-pol^/+* (*c(3)G^+^*) or *y w/ +; ru^1^ hry^1^ Diap1^1^(th) st^1^ cu^1^ c(3)G^mau^ /c(3)G^mau^;* sv*^spa-pol^/+* (*c(3)G^mau^*) females were mated individually to *ru^1^ hry^1^ Diap1^1^(th) st^1^ cu^1^ sr^1^ e^s^ ca^1^* males. Only female progeny were scored for the markers *ru hry Diap1^1^ (th) st cu*.

### Meiotic nondisjunction

To measure the rate of chromosome nondisjunction for both the *X* and *4th* chromosomes, virgin females of the indicated genotypes were mated to multiple *X^Y*, *In(1)EN*, *v f B; C(4)RM*, *ci ey^R^* males. The calculations to determine the percentage of *X* and *4th* chromosome nondisjunction, as well as adjusted progeny total, are described in (Hawley *et al*. 1992; Zeng *et al*. 2010; Zitron and Hawley 1989). Statistics test is described in (Zeng *et al*. 2010).

### Statistics and tetrad calculations

Fisher’s exact test was used to compare number of NCO/SCO to DCO/TCO numbers between genotypes and the difference for each interval in recombination experiments (https://www.graphpad.com/quickcalcs/contingency1/). Mann-Whitney U test was used to compare genotypes for centromere clustering, γH2Av foci number, and the distance between the mau C(3)G fluorescence (https://miniwebtool.com/mann-whitney-u-test-calculator/). The maximum likelihood estimates (MLE) of tetrad classes were calculated using the shiny app https://simrcompbio.shinyapps.io/CrossoverMLE/ (Hawley *et al*. 2025).

### Analysis of SC assembly and disassembly and γH2Av dynamics

Germaria stained with antibodies for mau C(3)G (SC), γH2Av (DSBs), and Orb (oocyte development) were analyzed for DSB and SC dynamics. For region 2A, only SC positive nuclei from cysts with strong Orb expression throughout the cyst were scored. Nuclei from different parts of 2A were scored, and typically more than one region 2A nucleus was analyzed for each germarium. For region 2B, if Orb had accumulated around a single nucleus in a cyst, that was the nucleus scored. If more than one cyst was present in region 2B, a nucleus was scored for DSB number from both cysts if the nuclei were unobstructed. For region 3, the Orb selected nucleus was scored for DSBs. To score DSB number, the nuclei of interest was duplicated in X, Y, and Z in Fiji and the individual Z slices made into a montage using a Make Montage Plugin. The γH2Av foci were then counted as they appeared through the Z slices.

For SC assembly state, the scored nuclei were described as one of four categories: 1) full-length with long, continuous tracks of SC; 2) mild fragmentation with some long tracks of SC and some partial tracks of SC; 3) highly fragmented where C(3)G is present but no long tracks of SC were observed; and 4) absent, in that there is a complete lack of SC structures in Orb selected nuclei (no germaria fell into this category). Only one value was assigned for each region of each germarium for SC assembly/disassembly. In region 2B, if the two cysts had differing SC classifications, the classification assigned was the more mature region 2B cyst.

## Results

*D. melanogaster* and *D. mauritiana* display a difference in crossover number and patterning despite their relatively close evolutionary history, similar genome sizes and similar amounts of pericentric heterochromatin (Hawley *et al*. 2025; True *et al*. 1996). *D. mauritiana* has an increased total map distance of 1.8-fold (463 vs. 259 cM) according to True *et al*. (1996) and 1.37-fold (379 vs 280 cM) for Hawley *et al*. (2025) compared to *D. melanogaster*. In both studies, the difference between *D. mauritiana* and *D. melanogaster* was not uniform between chromosome arms. Rather, the difference in total map distance is mostly explained by *D. mauritiana* having a decreased centromere effect compared to *D. melanogaster* which leads to greater genetic lengths on the chromosome arms that are amendable to crossover formation. While it has been shown previously that a transgenic *D. mauritiana mei-218* gene increased crossing over in *D. melangaster* (Brand *et al*. 2018), we believed additional trans-acting factors could also be influencing the difference in crossover patterning beteween the two species.

Prior studies in *D. melanogaster*, *C. elegans*, *A. thaliana,* and *Oryza sativa* ssp. *japonica* demonstrated that partial or complete loss of the SC can influence crossover placement (Billmyre *et al*. 2019; Capilla-Perez *et al*. 2021; Libuda *et al*. 2013; Wang *et al*. 2015). Additionally, current models propose that pro-crossover factors diffuse within the SC, and DSBs are selected to become crossovers through accumulation of pro-crossover factors by a coarsening process (Durand *et al*. 2022; Girard *et al*. 2023; Von Diezmann *et al*. 2024), indicating that changes in SC composition have the potential to alter coarsening dynamics, and ultimately, the number and position of crossovers.

By electron microscopy the SC looks like a ladder-like structure. It has a tripartite structure with two lateral elements (LEs) that interact with the homologous chromosomes. The distance between the LEs is spanned by transverse filament proteins that are stabilized by additional proteins within the central region of the SC. Fig. 1a shows the current model of the *D. melanogaster* SC. The *D. melanogaster* transverse filament protein C(3)G is modeled to be a homodimer with the C-termini of the homodimers interacting with the LEs (Anderson *et al*. 2005; Cahoon *et al*. 2017; Page and Hawley 2001). The N-termini of the C(3)G dimers interacts with homodimers from the opposing LE at the central element of the SC. Central region proteins Corona (Cona) and Corolla are thought to stabilize C(3)G assembly since absence of any of the three proteins result in a failure to assemble SC (Collins *et al*. 2014; Page and Hawley 2001; Page *et al*. 2008). As the transverse filament protein C(3)G is the major determinant of SC width (Anderson *et al*. 2005; Billmyre *et al*. 2019), this protein has a great potential for influencing crossover patterning. The C(3)G proteins in *D. mauritiana* and *D. melanogaster* share 89.1% identity with the amino acid changes dispersed throughout the protein (Supplementary Fig. 1 and 2). The level of amino acid difference between the species suggests the *D. mauritiana* C(3)G could potentially be structurally similar enough to interact with the other *D. melanogaster* proteins of the SC to allow for proper SC assembly but has enough amino acid difference that subtle differences in SC function might be found. To directly test whether C(3)G plays a role in the crossover patterning difference between the two species, the gene body of the *D. melanogaster c(3)G* gene was replaced with a codon-optimized coding region of the *D. mauritiana c(3)G* gene by WellGenetics using CRISPR-Cas9 technology (see Methods). This replacement is termed *c(3)G^mau^*. In the *c(3)G^mau^* flies, all other genes encoding central region and LE components of the SC still encode the endogenous *D. melanogaster* versions.

With the sequence divergence of C(3)G between the two species, there was the possibility that the C(3)G^mau^ protein would not be able to fully function within the context of building an SC structure with the *D. melanogaster* versions of the other components of the SC. We first examined SC assembly by immunofluorescence in *c(3)G^mau^* ovaries compared to the control line of similar genetic background, but with the endogenous *D. melanogaster c(3)G* (*c(3)G*^+^). In *D. melanogaster* wild-type ovaries, central region components of the SC first load near the centromeres during the mitotic divisions to form the 16-cell interconnected cyst (Fig. 1b)(reviewed in (Hughes *et al*. 2018)). As the cyst enters meiosis, SC components load at additional sites along the chromosome arms (zygotene). By region 2A (early pachytene), multiple nuclei of the cyst have formed full-length SC along the length of the chromosomes. As the cyst progresses to region 2B (early-mid pachytene), the SC progressively begins to disassemble from all but the nucleus selected to become the pro-oocyte, resulting in one nucleus with full-length SC by region 3 (mid-pachytene). After the cysts buds off to form the egg chamber, the SC in the pro-oocyte progressively disassembles (Wesley *et al*. 2020).

Based on an antibody recognizing the central region protein Corolla, SC assembly in regions 2A and 2B (early to early-mid pachytene) in *c(3)G^mau^* ovaries looked highly similar to *c(3)G^+^* with long continuous tracts of SC in numerous nuclei, indicating the C(3)G^mau^ protein could support full-length SC assembly with the other *D. melanogaster* SC proteins (Fig. 2). The monoclonal antibody raised against the C-terminus of *D. melanogaster* C(3)G fails to recognize the *D. mauritiana* C(3)G, so a polyclonal antibody raised against 324 amino acids in the C-terminal half of the *D. mauritiana* C(3)G protein (mau C(3)G) was generated (see Methods). The mau C(3)G antibody displayed a localization pattern similar to Corolla in the *c(3)G^mau^* replacement line, and it localized like Corolla in *c(3)G^+^* germaria expressing *D. melanogaster* C(3)G (Fig. 2). This indicates that the mau C(3)G antibody recognizes the C(3)G protein from both species.

To determine if the SC in *c(3)G^mau^* germaria has a tripartite structure, the localization of Corolla and mau C(3)G antibodies were examined by stimulated emission depletion (STED) microscopy. In both *c(3)G^+^* and *c(3)G^mau^* germaria, the fluorescence for the mau C(3)G antibody could be observed to split into two parallel tracks as has been observed previously by super-resolution microscopy for the *D. melanogaster* C-terminal antibody in wild type ovaries (Fig. 3)(Collins *et al*. 2014). In both *c(3)G^+^* and *c(3)G^mau^*, Corolla was located between the two tracks of C(3)G fluorescence, indicating that *c(3)G^mau^* assembles the expected tripartite SC structure. The distance between the two tracks of C(3)G fluorescence was measured to determine if the dimensions of the SC were altered. The average distance between the tracks of mau C(3)G antibody fluorescence in *c(3)G^mau^* was 112.1 nm (SD +/- 8.7 nm, N=25), while the distance between the mau C(3)G antibody staining in *c(3)G^+^* ovaries was significantly larger at 127.0 nm (SD +/- 11.6, N=27; p<0.0001). The tripartite pattern and continuous tracks in super-resolution microscopy indicates the C(3)G^mau^ protein can function with the *D. melanogaster* LE and central element proteins to promote tripartite SC assembly. The amino acid differences in the C(3)G^mau^ protein do not suggest an obvious explanation in the change in width compared to SC in *c(3)G^+^* ovaries. As 112.1 nm is within the width 90-150 nm range cited for SC in other organisms, it is unclear what functional consequence, if any, this difference in SC width would have on the functions of the SC.

**Figure 3.**
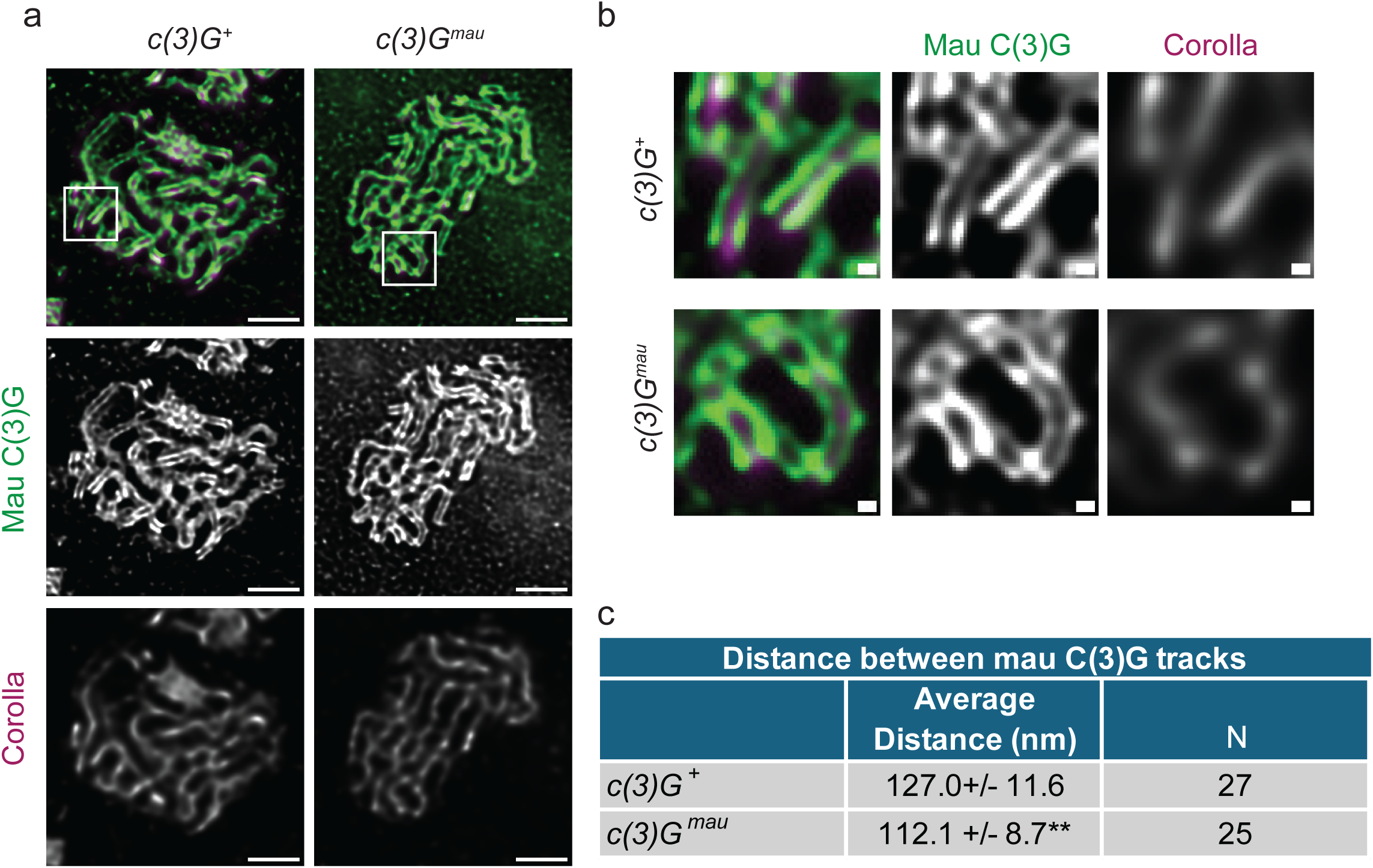
The SC in *c(3)G^mau^*ovaries has a tripartite structure but is narrower than SC from *c(3)G^+^* ovaries. a) Nuclei from *c(3)G^+^* and *c(3)G^mau^*acquired using STED microscopy. The mau C(3)G antibody against the C-terminal part of the protein can be resolved as two tracks (green) in both genotypes with an antibody to the central element protein Corolla localized between the tracks (magenta). Scale bar= 1 µm. Boxed region is magnified in b). Scale bar = 0.1 µm. c) Average and standard deviation are presented for measurements taken between the two tracks using the mau c(3)G antibody. N is the number of measurements. Statistically significant from *c(3)G^+^*at **p<0.01 by Mann-Whitney U test.

While tripartite SC is assembled in early pachytene in *c(3)G^mau^*ovaries, the stability of the full-length SC could be affected. We next examined SC dynamics within the germarium and observed full-length SC for the entirety of region 2A (early pachytene) where DSBs are normally initiated in *c(3)G^mau^* ovaries (Supplementary Fig. 3). The SC continued to appear full-length in the majority of germaria in region 2B (early-mid pachytene), which is the region DSBs continue to undergo repair and crossovers become designated based on previous data (Lake *et al*. 2015; Mehrotra and Mckim 2006)). The *c(3)G^+^* retained full-length SC in the nucleus selected to become the oocyte at region 3 (mid pachytene), consistent with previous studies (Supplementary Fig. 3) (Billmyre *et al*. 2019; Wesley *et al*. 2020). In *c(3)G^mau^*germaria, a mild early SC disassembly phenotype was observed. In region 3 approximately a quarter of the oocyte nuclei in *c(3)G^mau^* displayed some discontinuities in the tracks of SC (Supplementary Fig. 3). The fragmentation phenotype was mild in that most of these nuclei displayed a combination of long tracks of SC and some SC fragments. This mild onset of SC disassembly indicates the *C(3)G^mau^* containing SC may be less stable or its disassembly is being regulated differently compared to SC in *c(3)G^+^* germaria (Supplementary Fig. 3). Based on cytology, C(3)G^mau^ can function well enough with the *D. melanogaster* LE and central element components to support the formation of full-length tripartite SC in early pachytene despite the amino acid differences. The mild early SC disassembly phenotype suggests the amino acid changes in C(3)G^mau^ may lead to a protein that functions less well in maintaining SC than its *D. melanogaster* counterpart.

### C(3)G^mau^ protein supports wild-type levels of centromere clustering and chromosome segregation

We next examined the ability of C(3)G^mau^ to function in processes that depend on SC proteins. Studies in *D. melanogaster* have demonstrated that SC components are required for the clustering of homologous centromeres during pachytene into an average of two foci by early pachytene as based on immunofluorescence studies with an antibody recognizing the *Drosophila* homolog of CENPA, Centromere Identifier (CID)(Takeo *et al*. 2011; Tanneti *et al*. 2011). Null mutations of SC central region components in *D. melanogaster* result in a failure in centromere clustering (Collins *et al*. 2014; Takeo *et al*. 2011). The number of CID foci was scored along with antibodies against the SC (mau C(3)G) to identify meiotic nuclei and lamin to mark the nuclear boundary in both *c(3)G^+^* and *c(3)G^mau^* ovaries at different meiotic stages. An average of approximately two centromere clusters were observed at all meiotic stages examined for the *c(3)G^mau^* nuclei, which was not statistically significantly different from *c(3)G^+^* ovaries at corresponding stages, indicating the C(3)G^mau^ can function properly to cluster centromeres (Supplementary Fig. 4).

As C(3)G is required for the formation of the crossovers that ensure chromosome segregation at the meiosis I division (Page and Hawley 2001), we used a genetic assay to examine the segregation of the *X* and *4^th^*chromosomes in *c(3)G^mau^* females (Table 1). In assays of chromosome segregation, the *c(3)G^mau^* females displayed 0.2% *X* and 0.0% *4^th^* chromosome nondisjunction compared to 0.6% and 0.2% *X* and *4^th^* chromosome nondisjunction for *c(3)G^+^* females (Table 1). These levels of nondisjunction are not significantly different and thus, we conclude that sufficient crossovers are being generated to ensure homologous meiosis I chromosome segregation in *c(3)G^mau^* females. These results demonstrate the C(3)G^mau^ protein can substitute for the *D. melanogaster* version of C(3)G in the functions of centromere clustering and meiosis I chromosome segregation.

**Table 1.**
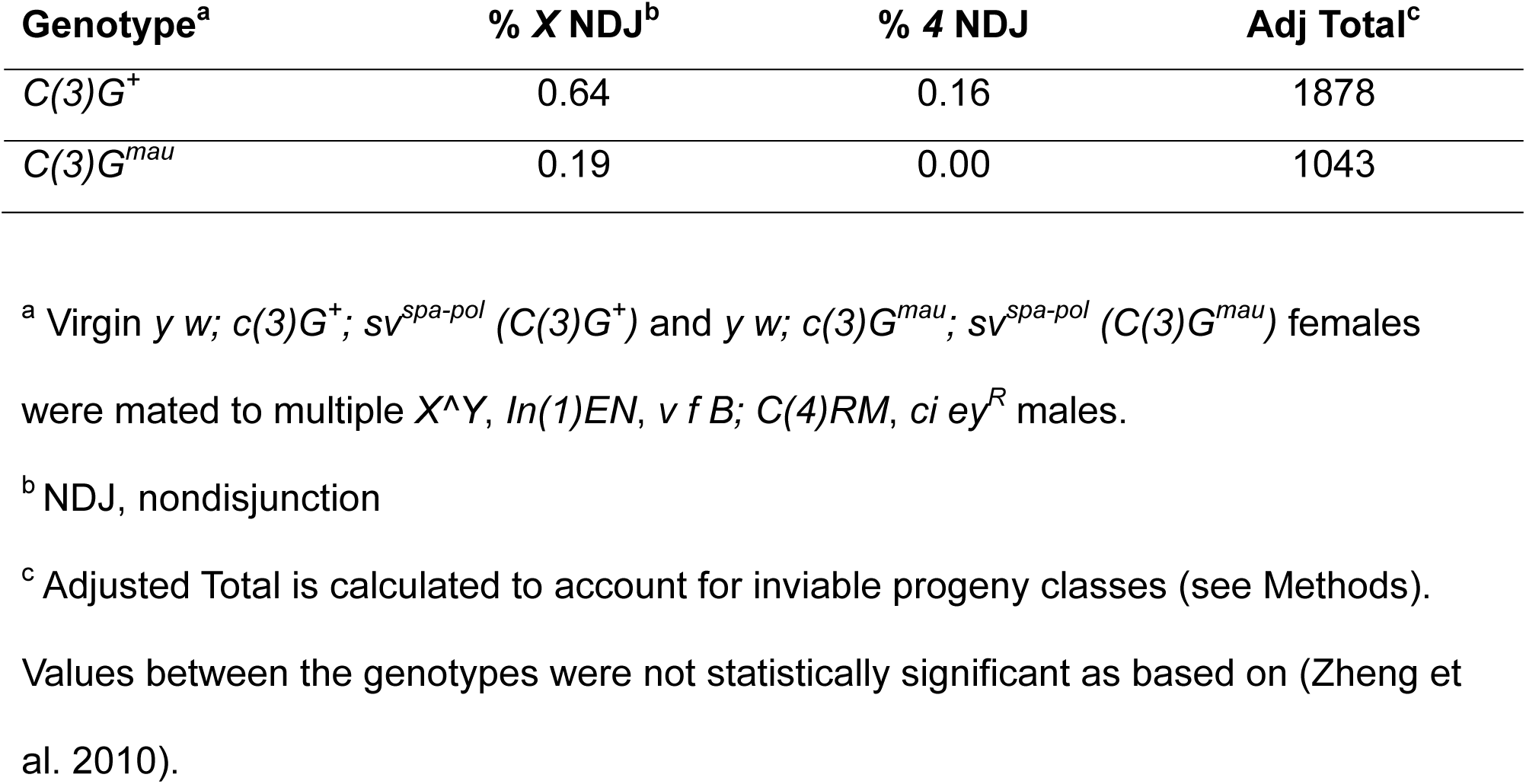
*X* and *4^th^* chromosome nondisjunction frequency.

### DSBs are induced at the correct stage

The SC has been shown to be necessary not only for crossover formation but also for the induction of wild-type levels of DSBs in early pachytene in *D. melanogaster* females (Collins *et al*. 2014; Mehrotra and Mckim 2006) (Fig. 1b). The induction and repair of DSBs were examined using antibodies recognizing γH2Av (DSBs), mau C(3)G (SC), and Orb (a marker of cyst development and oocyte selection). Numerous DSBs foci could be observed in *c(3)G^+^* and *c(3)G^mau^* ovaries in region 2A in cysts displaying strong Orb expression (*c(3)G^+^* 12.8 SD +/-2.5 foci compared to *c(3)G^mau^* 14.0 SD +/-4.6 foci) (Supplementary Fig. 5). In both genotypes at region 2B, the average number of γH2Av foci was reduced, indicating DSB repair is properly initiated in *c(3)G^mau^* ovaries.

The *c(3)G^mau^* ovaries did display an increase in the number of region 2B γH2Av foci (5.1 SD +/- 5.5 foci) compared to the *c(3)G^+^* control (0.8 SD +/- 2.1 foci) that was statistically different (p=0.0002). However, this number of region 2B foci in the *c(3)G^+^* ovaries was less than previously reported wild-type values for region 2B (Lake *et al*. 2013; Mehrotra and Mckim 2006). The difference in number of DSB foci at region 2B likely does not reflect a defect in DSB repair in the *c(3)G^mau^*ovaries since both genotypes displayed less than an average of one focus of γH2Av in region 3 (Supplementary Fig. 5). The small increase in DSB foci in region 2A and 2B may reflect a subtle effect on the kinetics of DSB repair due to the C(3)G^mau^ protein or potentially small differences in the kinetics of cyst development in the newly constructed *c(3)G^mau^* stock. The fact that *c(3)G^mau^*females displayed wild-type levels of chromosome segregation (Table 1) further suggests that that the small differences in DSB kinetics are not an indication of defects in the proper repair of DSBs. The use of the Orb antibody for the DSB assay also allowed for the examination of the degree of activation of the pachytene checkpoint, a meiotic delay that can be induced by defects in the chromosome axis or DSB repair (Joyce and Mckim 2009; Joyce and Mckim 2010). Activation of the pachytene checkpoint leads to a persistence of strong Orb accumulation in the cytoplasm around two nuclei in region 3, the “two-oocyte phenotype”(Joyce and Mckim 2009; Joyce and Mckim 2010). Out of 21 *c(3)G^mau^* germaria, 20 displayed a single nucleus with Orb concentrated in the cytoplasm with the remaining germarium displaying a nucleus with strong Orb accumulation in the cytoplasm and a second nucleus with intermediate Orb concentration. In all 18 of the *c(3)G^+^* ovaries examined, Orb was concentrated around a single nucleus at region 3. This result suggests that there is not a strong induction of the pachytene checkpoint in *c(3)G^mau^* ovaries. Overall, these data indicate that *C(3)G^mau^*can function with the *D. melanogaster* SC components to promote induction of wild-type levels of DSBs at early pachytene and the repair of DSBs by region 3 is sufficient to the point that γH2AV is absent in the selected oocyte nucleus.

### Crossover patterning is altered in *c(3)G^mau^* females and appears more similar to D. mauritiana *than* D. melanogaster

The chromosome segregation assays described above indicate that a sufficient level of crossovers are formed for proper chromosome segregation. However, we wanted to determine if the number and placement of crossovers were affected by the utilization of the C(3)G^mau^ protein. Recombination along the entire *X* chromosome was assayed in both *c(3)G^+^* and *c(3)G^mau^* females by scoring visible markers on a multiply marked *X* chromosome. First, we examined the distribution of the types of recovered crossover chromatids. There was a decrease in the portion of non-crossover (NCO) chromatids recovered from *c(3)G^mau^* compared to *c(3)G^+^* females (36.5% compared to 47.0%)(Fig. 4, Table 2). This decrease in NCO frequency was accompanied by a proportional increase in the number of double and triple crossover chromatids (DCO and TCO) recovered from *c(3)G^mau^*females. The fraction of single crossover chromatids (SCO) between the two genotypes was similar (Table 2). The difference between the proportion of NCO/SCO to DCO/TCO classes between *c(3)G^mau^* and *c(3)G^+^* was statistically significant (Fisher exact test table p<0.0001), indicating an excess of chromosomes experiencing more than one crossover.

**Figure 4.**
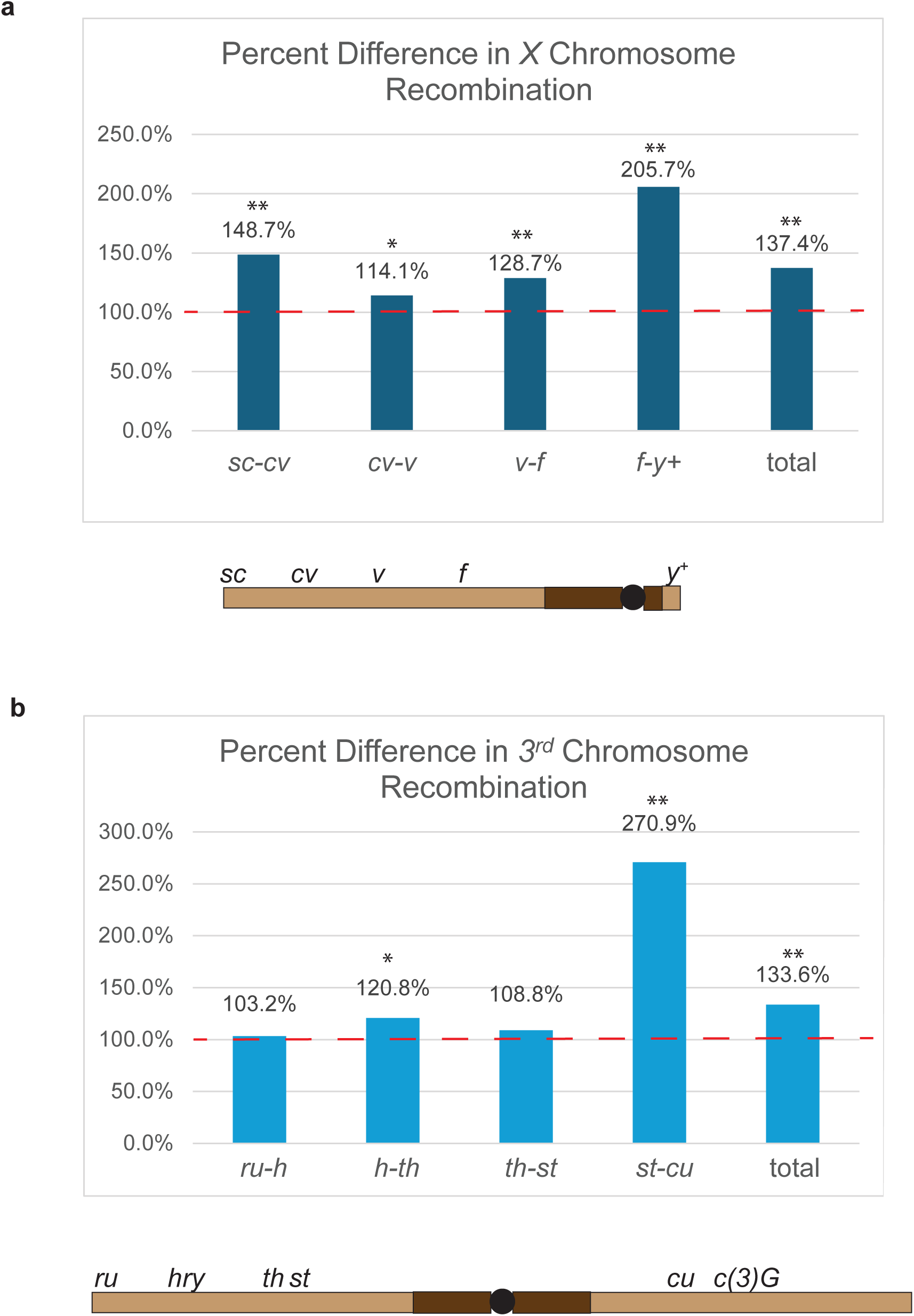
Recombination in the interval spanning the centromere is increased in *c(3)G^mau^* females. a) The percent difference in *X* chromosome recombination observed for *c(3)G^mau^* compared to the *c(3)G^+^*control (Red dotted line set to 100% of control) for the indicted intervals. Arrangement of the markers scored on the X chromosome are diagramed below the graph. b) The percent difference in *3^rd^* chromosome recombination observed for *c(3)G^mau^* compared to the *c(3)G^+^* control (Red dotted line set to 100% of control) for the indicted intervals. Arrangement of the markers scored on the *3rd* chromosome, as well as the location of *c(3)G*, are diagramed below the graph. Statistically significant from *c(3)G^+^* at * p<0.05 and **p<0.01.

**Table 2.**
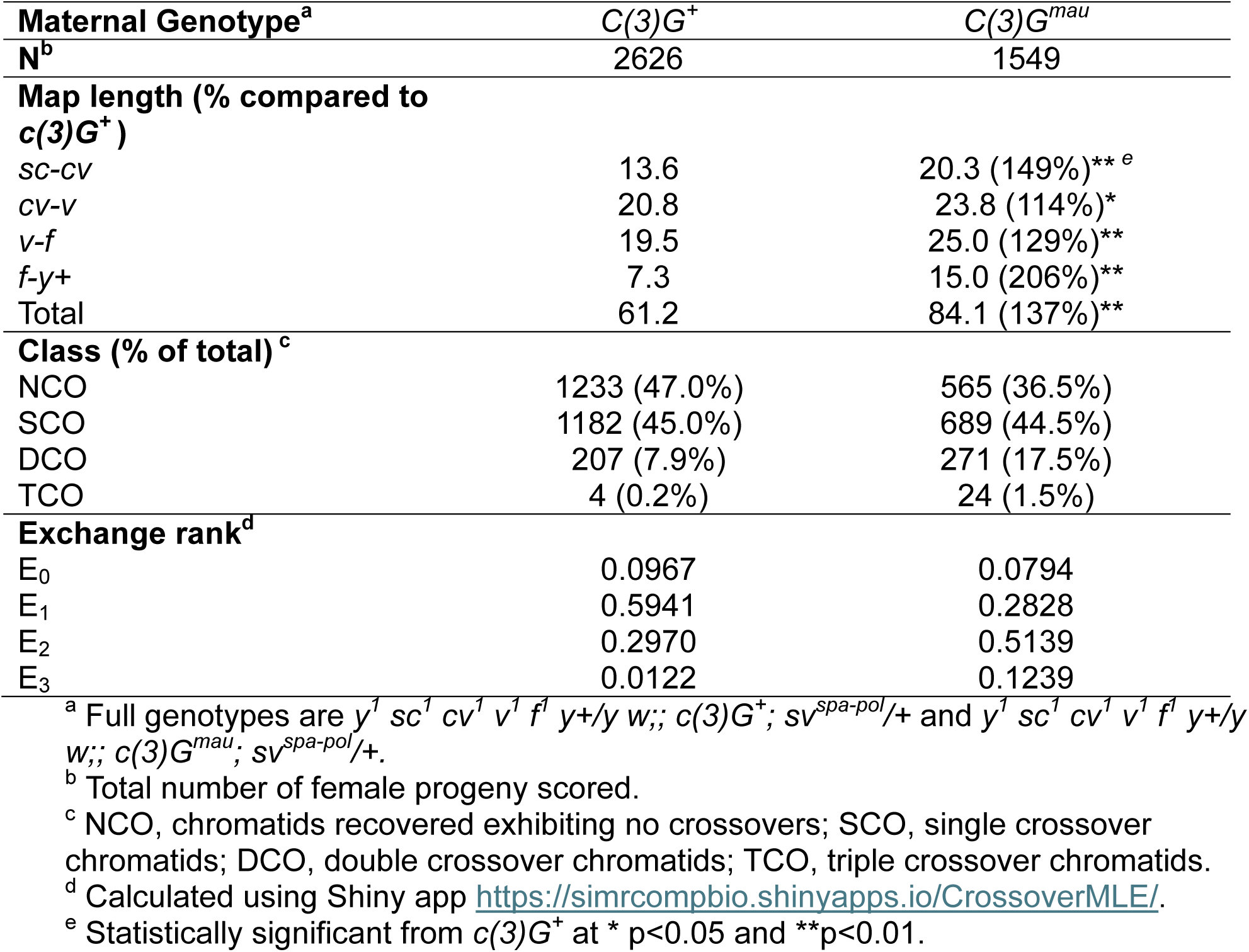
*X* Chromosome Recombination.

The rate of crossing over within each interval of the *X* chromosome was compared between *c(3)G^+^* and *c(3)G^mau^,* revealing increases in crossing over in *c(3)G^mau^* females for all scored intervals (Fig. 4, Table 2). Fig. 4 displays the recombination rate of *c(3)G^mau^* as a percentage of *c(3)G^+^*to illustrate the difference in the recombination pattern between the two genotypes. While every interval examined showed a statistically significant increase in crossing over for the *c(3)G^mau^* females, the most striking increase was in the interval spanning the centromere between *forked (f)* and the wild-type *yellow (y+)* marker translocated to the small right arm of the *X* chromosome (Fig. 4, Table 2), which is the most centromere-proximal interval we examined. Crossing over was increased by 206% in *c(3)G^mau^*females compared to *c(3)G^+^* females, indicating that recombination most substantially increased in the centromere-proximal euchromatin of the *X* chromosome. This increase in crossing over in centromere-proximal euchromatin mirrors the strong increase in recombination observed in the centromere-proximal euchromatin of the *D. mauritiana* species (Hawley *et al*. 2025; True *et al*. 1996).

To further examine the level of crossing over, Weinstein Exchange Ranks were calculated using Weinstein tetrad analysis for *c(3)G^mau^* and *c(3)G^+^* (Hawley *et al*. 2025; Weinstein 1936). As only one of the four chromatids can be recovered in the progeny during female meiosis, the actual proportion of chromatids with 0,1,2, or more crossovers must be mathematically derived from the recovered progeny and these exchange classes are classified as E0, E1, E2, and E3 respectively. (https://simrcompbio.shinyapps.io/CrossoverMLE/ )(Hawley *et al*. 2025). For the *X* chromosome the estimated E0 frequency for *c(3)G^mau^* was 0.0793 compared to 0.0967 for *c(3)G^+^*(Table 2). The biggest shift was in the calculated E2 class with values of 0.5139 for *c(3)G^mau^*and 0.297 for *c(3)G^+^*. The values for *c(3)G^mau^*females share similarities with Weinstein tetrad frequencies for the *X* chromosome observed in whole-genome sequencing studies in *D. mauritiana* (Hawley *et al*. 2025). The estimated *X* chromosome E0, E1, and E2 values for *D. mauritiana* from that study were 0.1048, 0.3238, and 0.5714, respectively (Hawley *et al*. 2025). The *X* chromosome Weinstein tetrad frequencies estimated from previous *D. melanogaster* whole-genome sequencing data were 0.1224, 0.6122, and 0.2654 for E0, E1, and E2, respectively (Hawley *et al*. 2025; Miller *et al*. 2016). The *c(3)G^+^*control matches well with the whole genome sequencing data of *D. melanogaster* while the *c(3)G^mau^* results are much more similar to the values observed for *D. mauritiana* (Hawley *et al*. 2025; Miller *et al*. 2016). The Weinstein tetrad frequencies further illustrate the increase in crossing over in *c(3)G^mau^* females and that this increase has similarities to the crossover pattern of *D. mauritiana* females for the *X* chromosome.

Previous studies examining *D. melanogaster* females with internal deletion mutations in the C(3)G protein have demonstrated that the recombination pattern of the *X* chromosome and an autosome can be altered in differing ways (Billmyre *et al*. 2019).

Additionally, while the *D. mauritiana* species showed increases in crossing over and a decreased centromere effect compared to *D. melanogaster*, the magnitude of those differences were not uniform across all five chromosome arms that experience crossing over (Hawley *et al*. 2025; True *et al*. 1996). We next examined crossing over on the *3^rd^* chromosome from the marker *roughoid* (*ru*) near the telomere of *3L* to *curled* (*cu*) which is located across the *3^rd^* chromosome centromere on *3R*. Like the *X* chromosome, there was an increase in DCO chromatids (10.6% for *c(3)G^mau^* compared to 5.2% for *c(3)G^+^*) at the expense of NCO chromatids in *c(3)G^mau^* (41.7%) compared to *c(3)G^+^* (53.8%)(Table 3). As seen for the *X* chromosome, the difference in the proportion of NCO/SCO chromatids compared to DCO/TCO chromatids was statistically significant for *c(3)G^mau^* compared to *c(3)G^+^* females (Fisher exact test table p<0.0001).

**Table 3.** *3rd* Chromosome Recombination.

| Maternal Genotype<sup>a</sup> | C(3)G<sup>+</sup> | C(3)G<sup>mau</sup> |
| --- | --- | --- |
| N<sup>b</sup> | 1862 | 1589 |
| **Map length (% compared to c(3)G<sup>mel</sup>)** |  |  |
| *ru-h* | 24.8 | 25.6 (103%)<sup>e</sup> |
| *h-th* | 19.1 | 23.0 (121%)* |
| *th-st* | 0.8 | 0.8 (109%) |
| *st-cu* | 7.4 | 20.1 (271%)** |
| Total | 52.0 | 69.5 (134%)** |
| **Class (% of total)<sup>c</sup>** |  |  |
| NCO | 1001 (53.8%) | 662 (41.7%) |
| SCO | 759 (40.8%) | 753 (47.4%) |
| DCO | 96 (5.2%) | 169 (10.6%) |
| TCO | 6 (0.3%) | 5 (0.3%) |
| **Exchange rank<sup>d</sup>** |  |  |
| E<sub>0</sub> | 0.178 | 0.046 |
| E<sub>1</sub> | 0.628 | 0.541 |
| E<sub>2</sub> | 0.168 | 0.388 |
| E<sub>3</sub> | 0.026 | 0.025 |

<sup>a</sup> Full genotypes are *y w/+;; ru<sup>1</sup> hry<sup>1</sup> Diap1<sup>1</sup>(th) st<sup>1</sup> cu<sup>1</sup> sr<sup>1</sup> e<sup>s</sup> ca<sup>1</sup> /+; sv<sup>spa-pol</sup> /+* and *y w/+;; ru<sup>1</sup> hry<sup>1</sup> Diap1<sup>1</sup>(th) st<sup>1</sup> cu<sup>1</sup> c(3)G<sup>mau</sup> /c(3)G<sup>mau</sup>; sv<sup>spa-pol</sup> /+*.
<sup>b</sup> Total number of female progeny scored.
<sup>c</sup> NCO, chromatids recovered exhibiting no crossovers; SCO, single crossover chromatids; DCO, double crossover chromatids; TCO, triple crossover chromatids.
<sup>d</sup> Calculated using the Shinyapp at https://simrcompbio.shinyapps.io/CrossoverMLE/
<sup>e</sup> Statistically significant from c(3)G<sup>+</sup> at \* p<0.05 and \*\*p<0.01.

Finally, the level of recombination within each of the tested intervals was examined. For the most centromere distal interval, *ru-h,* and the small *Diap1(th)-st* interval recombination levels were similar between *c(3)G^mau^* and *c(3)G^+^* (Fig. 4, Table 3). A small, but statistically significant, increase in recombination was observed for the *h-Diap1(th)* interval in the medial part of the *3L* chromosome arm for *c(3)G^mau^* compared to *c(3)G^+^* females (Fig. 4, Table 3). What was most noticeable was the 271% increase in crossing over in the *st-cu* interval for *c(3)G^mau^* compared to *c(3)G^+^* (Fig. 4, Table 3). This interval encompasses the centromere and the centromere-proximal euchromatin on both sides of the centromere (Fig. 4). This strong increase in recombination in the centromere-spanning intervals on both the *X* and *3^rd^* chromosomes suggests a general weakening of the spread of the centromere effect into the centromere-proximal euchromatin in the *c(3)G^mau^* females. In our recent recombination study on *D. mauritiana*, two of the regions with the most significant increase in crossing over compared to *D. melanogaster* were the centromere-proximal regions of the *3^rd^* chromosome (Hawley *et al*. 2025).

To better evaluate the decrease in NCO chromatids, Exchange Ranks were again calculated using Weinstein tetrad analysis. Similar to the *X* chromosome, *c(3)G^mau^* females displayed a decrease in the calculated E0 frequency and an increase in E2 values compared to the control (*c(3)G^mau^*E0=0.0459 and E2=0.3877 compared to *c(3)G^+^* E0=0.1783 and E2=0.1676) (Table 3). Altogether, our results indicate that the presence of C(3)G^mau^ protein in *D. melanogaster* females is sufficient to increase crossing over on multiple chromosome arms and generate a crossover pattern more similar to the *D. mauritiana* species than wild-type *D. melanogaster* females.

## Discussion

### *Cis* versus *trans-*acting factors regulating changes in crossover patterning

The number of crossovers formed per bivalent and their location on the chromosomes can vary between species, leading to species-specific crossover patterns that can differ even between closely related species. It is not fully understood how crossover patterns can evolve. Are new crossover patterns driven by changes in just a few genes or are crossover patterns derived from the small effects of many genes and cis-acting genetic elements?

Here, we show that complete replacement of the *D. melanagaster c(3)G* gene with *D. mauritiana c(3)G* alters the recombination pattern of *D. melanogaster* to look more like the *D. mauritiana* crossover pattern, with increased crossing over in the centromere-proximal euchromatin and an increase in total crossing over for the *X* and *3^rd^* chromosomes. Examining the calculated E0 frequencies from these experiments revealed a decrease in NCO chromatids in *c(3)G^mau^*females. As *c(3)G^mau^*females displayed wild-type levels of chromosome segregation, it is clear that the changes observed in crossover placement and number were not detrimental to this critical function of meiosis. Given that this substitution represents a single gene, this result suggests that global crossover patterning can be strongly influenced by a relatively small number of genetic loci. Additionally, this result supports the inference that the C(3)G protein and/or the SC is a mechanism to regulate crossover patterning in *Drosophila*. *c(3)G* is not the first example of a single gene noticeably altering crossover patterning in *D. melanogaster*. Brand *et al*. (2018) expressed the *D. mauritiana* version of the pro-crossover protein Mei-218 as a transgene in *D. melanogaster* that lacked a functional endogenous *mei-218* gene. Expression of *D. mauritiana mei-218* increased total crossing over, particularly for genetic intervals near the telomere and centromeres. However, in that study, it was not apparent that a wild-type genetic map could be fully restored, with either the *D. mauritiana* or *D. melanogaster* transgenes. Here, with the full replacement of the *D. melanogaster c(3)G* with that of *D. mauritiana,* a genetic map quite similar to *D. mauritiana* was observed. This result provides another example of how a single gene can influence both the placement and rate of crossovers. In comparison to the *mei-218* transgene results, the *c(3)G^mau^* replacement had a lesser impact on regions near the telomere with the increases in recombination most focused on intervals spanning the centromeres.

Unlike studies directly comparing recombination between the *D. melanogaster* and *D. mauritiana* species, where the centromere and centromere-proximal heterochromatin are different (Hawley *et al*. 2025; True *et al*. 1996), in the *mei-218* and *c(3)G* studies the *cis-*acting elements at or near the centromere should be essentially identical between the compared flies. Therefore, the changes observed in crossover patterning within the *D. melanogaster* females could only be driven by the *trans-*acting factors that were manipulated in the studies. This strongly suggests that the difference in the strength of the centromere effect between *D. melanogaster* and *D. mauritiana* is primarily mediated by *trans*-acting factors, rather than specific *cis*-acting elements within the centromeres and surrounding heterochromatin of those species. *Cis* acting elements, such as heterochromatin and hotspots, would still influence local placement of crossovers but these *trans*-acting factors apparently enable global changes in patterning that are observed across species.

Why were Mei-218 and C(3)G good candidates for regulating aspects of global crossover patterning? Both are rapidly evolving proteins (Brand *et al*. 2018; Hemmer and Blumenstiel 2016; Zakerzade *et al*. 2025) and are involved in the formation of crossovers in *D. melanogaster* (Mckim *et al*. 1996; Page and Hawley 2001). While genes involved in meiosis tend to show higher rates of evolution than genes that function in the soma, Mei-218 was chosen to test because it was found to be particularly fast evolving (Brand *et al*. 2018). While the rate of C(3)G divergence is less than Mei-218, C(3)G and other components of the SC are evolving so rapidly that they cannot readily be identified outside the genus based on amino acid sequence alone (Hemmer and Blumenstiel 2016; Zakerzade *et al*. 2025). The rapid divergence of these proteins makes both C(3)G and Mei-218 good candidates for driving rapid evolution of the recombination landscape. If evolution of the recombination landscape occurred only by *cis-*acting factors, rapid evolution of the global landscape would require many changes occurring in a short span of time, which seems less plausible.

### The role of the SC in crossover patterning

How might the SC alter recombination patterns? Current models predict that pro-crossover factors can diffuse within the SC and then, through a coarsening process, the pro-crossover factors accumulate at the subset of DSBs that become designated to become crossovers (Durand *et al*. 2022; Girard *et al*. 2023; Von Diezmann *et al*. 2024). It was recently shown that a mutation in the *C. elegans* SC component, *syp-4*, can promote SC assembly but also causes an increased number of crossovers (Kohler *et al*. 2025). Examination of the SC by three-dimensional stochastic optical reconstruction microscopy (3D-STORM) revealed changes in the organization of the components of the SC in the *syp-4* mutant (Kohler *et al*. 2025). Internal deletion mutations in c(3)G also cause defects in SC assembly and/or disassembly, as well as changes in crossover levels and placement (Billmyre *et al*. 2019). More recently, it was shown that there is an increase in crossovers in the β-heterochromatin associated with the *2^nd^*chromosome centromere in one of these deletion mutants, indicating C(3)G and/or the SC plays a role in suppressing crossovers within parts of the centromeric heterochromatin (Pazhayam *et al*. 2025).

Our work here demonstrated that the *C(3)G^mau^* replacement can promote full-length tripartite SC assembly, chromosome segregation, and centromere clustering, indicating this version of C(3)G can interact well enough with *melanogaster* meiotic components to promote these important functions. The initiation of premature SC disassembly was observed in about a quarter of germaria at region 3 (mid pachytene), but this phenotype is mild compared to the phenotype of *c(3)G* internal deletion mutants (Billmyre *et al*. 2019) and occurs after crossovers are designated based on the localization of the RING domain protein Vilya (Lake *et al*. 2015). The timing and mildness of the disassembly phenotype suggests the change in crossover patterning observed in *c(3)G^mau^* females cannot be simply explained by a loss of SC structure. While tripartite SC structure was observed by super-resolution microscopy in *c(3)G^mau^*, a statistically significant decrease in width between the tracks of mau C(3)G antibody fluorescence was also observed.

The difference in width may contribute to differences in SC function. Along with the mild disassembly phenotype, the difference in SC width opens the possibility there may be some difference in the organization of the SC components in *c(3)G^mau^* ovaries. If use of the C(3)G^mau^ protein caused changes in the interaction of SC proteins similar to the recently described *syp-4* mutant in *C. elegans* (Kohler *et al*. 2025), the altered SC structure could potentially alter how pro-crossover factors diffuse within the SC to ultimately affect the number and location of DSBs designated to become crossovers. In studies of complete reconstructions of electron microscopy (EM) data of nuclei from fly germaria Carpenter (1975) described the SC of the heterochromatin approaching the chromocenter as having a different morphology than the SC of euchromatin, with less distinct LEs and a relatively amorphous central element. The recent identification of crossovers in the β -heterochromatin in a *c(3)G* deletion mutant suggests that the morphological difference in SC structure could potentially be part of the mechanism of the centromere effect in *D. melanogaster,* perhaps by varying the manner in which centromeric suppression is transmitted along the chromosome arm. It would be interesting to use EM in the future to explore whether the C(3)G^mau^ protein could potentially be altering the organization of the SC in and near the centromere-proximal heterochromatin. One can imagine a change in SC structure in these regions could make recruitment of pro-crossover factors to the euchromatic regions closer to the centromeres more likely and increase crossing over in the centromere-proximal euchromatin.

The SC was observed in EM images decades earlier (Moses 1956), and its role in promoting crossover formation in many species has been of intense interest. Only more recently have there been studies on mutations or conditions that effect the SC without disrupting the structure entirely. These studies are beginning to reveal additional, influential roles of this conserved structure and are providing information on how crossover patterns may be regulated and altered by SC evolution.

## Data Availability Statement

Original data underlying this manuscript can be accessed from the Stowers Original Data Repository at http://www.stowers.org/research/publications/libpb-2555.

## Acknowledgements

Thank you to Cathy Lake, Jay Unruh, and Leah Rosin for valuable comments on the manuscript and Stefanie Williams for her help with figure preparation and suggestions.

## Funding

Funding for this work came from the Stowers Institute for Medical Research (R.S.H) and National Science Foundation award 2025197 (JB). R.S.H. was an American Cancer Society Research Professor.

**Supplementary Figure 1. Codon optimized *D. mauritiana c(3)G* coding sequence.** The codon optimized nucleotide *D. mauritiana c(3)G* coding sequence (line 1) is shown aligned with publicly available sequences for *D. mauritiana* (line 2) and *D. melanogaster c(3)G* coding sequences (line 3). Amino acid sequence coded by each codon is also displayed illustrating that the codon optimized nucleotide sequence retained the amino acid sequence of the publicly available *D. mauritiana* sequence.

**Supplementary Figure 2. Alignment of D. *melanogaster* and *D. mauritiana* C(3)G proteins.** Amino acid differences are displayed in red and gaps in white. The proteins show 89.1% identity with differences dispersed throughout the protein. The region expressed to generate the *D. mauritiana* C(3)G antibody is underlined.

**Supplementary Figure 3. SC fully assembles in region 2A (early pachytene) but begins to disassemble early in some c*(3)G^mau^* ovaries.** a) Examples of nuclei from *c(3)G^mau^* ovaries at region 3 (mid pachytene) with full length, mildly fragmented, or highly fragmented SC. SC was labeled with the mau C(3)G antibody. Images are projections from partial z-stacks and scale bar = 1 µm. b) Shown is the percentage of germaria with the scored SC assembly state for *c(3)G^+^* (N=18 germaria) and *c(3)G^mau^* (N=21 germaria) for germarium regions 2A (early pachytene), 2B (early-mid pachytene) and 3 (mid pachytene).

**Supplementary Figure 4. Centromeres cluster properly in c*(3)G^mau^* ovaries**. a) Examples of nuclei from *c(3)G^mau^*ovaries with 1 (top row), 2, or 3 (bottom row) centromere clusters. Centromeres are labeled with an antibody recognizing the centromeric histone CID (magenta), mau C(3)G antibody labels the SC (yellow), and lamin antibody outlines the nuclear envelope (cyan). Asterisks indicate the location of centromere clusters. Scale bar= 1 µm and images are projections from partial z-stacks. b) The average number of centromere clusters with standard deviation at the indicated developmental stages in *c(3)G^+^* and *c(3)G^mau^*ovaries. Number of nuclei scored in parentheses. Mann-Whitney U test found no statistically significant difference between the genotypes for any of the stages examined (p>0.05).

**Supplementary figure 5. Double strand breaks are induced at early pachytene and undergo repair in *c(3)G^mau^* ovaries.** a) *c(3)G^+^*(top) and *c(3)G^mau^* (bottom) germaria were stained with antibodies to mau C(3)G to identify the SC (green), γH2Av to recognize DSBs (magenta), Orb to stage oocyte development (white), and DAPI (blue). In both genotypes DSBs are induced in region 2A (early pachytene) and γH2Av staining is absent from the Orb selected nucleus in region 3. Note in the *c(3)G^mau^* image γH2Av foci are present in the nurse cells that have started endoreduplication in region 3. Scale bar = 10 µm and images are projections from z stacks. b) Table provides the average number with standard deviation of γH2Av foci for each region of the germarium. The total number of nuclei scored is in parentheses. By Mann-Whitney U test the statistically significant differences from *c(3)G^+^* are indicated as * p<0.05 and **p<0.01. Full genotypes are *y w; c(3)G^+^; spa^pol^*and *y w; c(3)G^mau^; spa^pol^*.

